# *C. elegans* MRP-5 Exports Vitamin B12 from Mother to Offspring to Support Embryonic Development

**DOI:** 10.1101/204735

**Authors:** Huimin Na, Olga Ponomarova, Gabrielle E. Giese, Albertha J.M. Walhout

## Abstract

Vitamin B12 functions as a cofactor for methionine synthase to produce the anabolic methyl donor S-adenosylmethionine (SAM) and for methylmalonyl-CoA mutase to catabolize the short chain fatty acid propionate. In the nematode *Caenorhabditis elegans*, maternally supplied vitamin B12 is required for the development of her offspring. However, the mechanism for exporting vitamin B12 from the mother to her offspring is not yet known. Here, we use RNAi of more than 200 transporters with a vitamin B12-sensor transgene to identify the ABC transporter MRP-5 as a candidate vitamin B12 exporter. We show that injection of vitamin B12 into the gonad of *mrp-5* deficient mothers rescues embryonic lethality in her offspring. Altogether, our findings identify a maternal mechanism for the transit of an essential vitamin to support the development of the next generation.

## INTRODUCTION

Maternal micronutrient status during pregnancy greatly affects embryonic development and fetal health (Dror and Allen, 2008; Fall et al., 2003; Owens and Fall, 2008; Skjaerven et al., 2016). For instance, the intake of folate supplements by pregnant women reduces the occurrence of neural tube defects in newborns (Czeizel and Dudas, 1992; Viswanathan et al., 2017). Folate functions together with vitamin B12, or cobalamin, in the one carbon cycle to recycle methionine from homocysteine (Moreno-Garcia et al., 2013)(**Figure 1A**). One of the main functions of the one-carbon cycle is to produce S-adenosylmethionine (SAM), a major methyl donor required to synthesize phosphatidylcholine, an important component of cellular membranes, and to modify histones and DNA to ensure proper genomic functions (Rosenblatt and Whitehead, 1999). The other function of vitamin B12 is as a cofactor in the breakdown of the short chain fatty acid propionate (Carrillo-Carrasco and Venditti, 1993)(**Figure 1A**). Humans obtain vitamin B12 through ingestion of animal products in which this stable micronutrient is efficiently stored (Martens et al., 2002). Deficiency of vitamin B12 in newborns is mostly caused either by maternal depletion or malabsorption, although rare recessive inborn errors of vitamin B12 processing or utilization have also been identified (Rosenblatt and Cooper, 1990; Rosenblatt and Whitehead, 1999).

**Figure 1.**
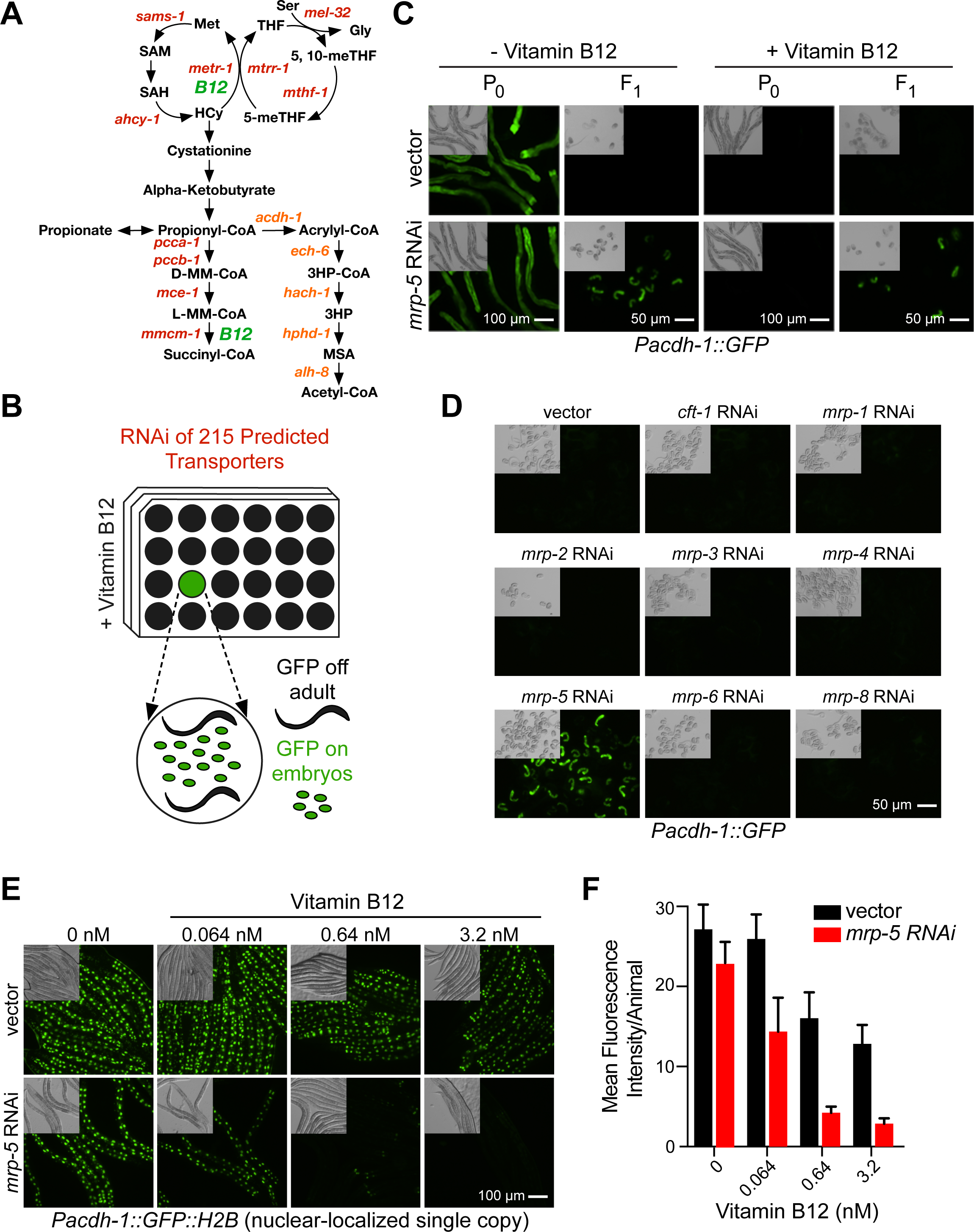
A *C. elegans* Transporter RNAi Screen Identifies *mrp-5* as a Potential Vitamin B12 Exporter. (A) Cartoon of the two vitamin B12-dependent metabolic pathways. Gene names are from *C. elegans*. 3HP – 3-hydroxypropionate; 5-meTHF −5-methyltetrahydrofolate; 5, 10-meTHF - 5,10-methylenetetrahydrofolate; D-MM-CoA – D-methylmalonyl-CoA; Gly – glycine; HCy – homocysteine; L-MM-CoA – L-methylmalonyl-CoA; Met – methionine; MSA – malonic semialdehyde; SAH – S-adenosyl homocysteine; SAM – S-adenosyl methionine; THF – tetrahydrofolate. The *acdh-1* branch in orange text indicates the vitamin B12-independent propionate shunt. (B) Diagram of *C. elegans* transporter RNAi library screen. RNAi-treated animals that displayed GFP fluorescence in the F1 but not the P0 generation in the presence of 64 nM vitamin B12 were considered hits. (C) Fluorescence and DIC microscopy images of *Pacdh-1*∷GFP animals subjected to RNAi of *mrp-5* in *E. coli* HT115 compared to vector control at P0 and F1 generations in the presence or absence of 64 nM vitamin B12. Scale bars, 50 μm for embryos and 100 μm for adults. (D) Fluorescence and DIC microscopy images of *Pacdh-1∷GFP* animals subjected to RNAi of *mrp* family members in *E. coli* HT115 compared to control RNAi in the F1 generation. An RNAi clone for *mrp-7* was not available. Scale bar, 50 μm. (E) Fluorescence microscopy images of single copy insertion *Pacdh-1∷GFP∷H2B* animals subjected to *mrp-5* RNAi in *E. coli* OP50 compared to control RNAi with low dose titration of vitamin B12. Scale bar, 100 μm. (F) Imagines from part E were quantitated using ImageJ to determine the mean fluorescence intensity per animal. At least five animals were analyzed per condition.

Recently, the nematode *Caenorhabditis elegans* has been established as a powerful model for studying vitamin B12-dependent processes (Yilmaz and Walhout, 2014). *C. elegans* is a bacterivore that can be fed individual bacterial species and strains that can elicit different effects on the animal’s life history traits and gene expression. For instance, when fed *Comamonas aquatica* DA1877 bacteria, the animals develop faster, have reduced fecundity, and shorter lifespan relative to animals fed the standard laboratory diet of *Escherichia coli* OP50 (MacNeil et al., 2013; Watson et al., 2013). These dietary differences are due, at least in part, to the fact that *Comamonas* synthesizes vitamin B12, whereas *E. coli* does not (Watson et al., 2014). Dietary vitamin B12 is required to support *C. elegans* development, as the animal itself cannot synthesize this cofactor (Bito et al., 2013; Watson et al., 2014). Vitamin B12 is ingested by the mother and needs to be exported from the intestine into the gonad to support the development of her offspring. However, the mechanism by which this occurs is not yet known. Here, we report that the ABC transporter *mrp-5* controls vitamin B12 export from the intestine to support *C. elegans* embryonic development.

## RESULTS

### An RNAi Screen Identifies the ABC Transporter MRP-5 as a Candidate to Export Vitamin B12 from the *C. elegans* Intestine

We have previously established the *Pacdh-1∷GFP* transgenic *C. elegans* strain as a reporter of dietary vitamin B12 status: in this strain GFP is highly expressed when vitamin B12 is low and repressed when levels of this micronutrient are high (Arda et al., 2010; MacNeil et al., 2013; Watson et al., 2014; Watson et al., 2016). GFP expression in this strain is under the control of the promoter of the acyl-CoA dehydrogenase *acdh-1*. This gene encodes the first enzyme of an alternate propionate breakdown pathway, or propionate shunt, which does not require vitamin B12 and is transcriptionally activated when this cofactor is in low supply (Watson et al., 2016)(**Figure 1A**). We reasoned that a defect in vitamin B12 transport from the mother to her offspring would result in P0 animals with low levels of GFP in the presence of vitamin B12 and F1 embryos with high levels of GFP expression. Further, these embryos would be expected to have developmental defects because vitamin B12 is required to produce SAM, which is needed to generate biomass and thus support development (Watson et al., 2014). We performed RNAi knockdown of 215 predicted *C. elegans* transporters of the ABC transporter and solute carrier families by feeding animals *E. coli* HT115 bacteria expressing double-stranded RNA in the presence of 64 nM vitamin B12 (**Figure 1B**). We identified a single gene, *mrp-5*, that, when knocked down, resulted in mothers with low levels of GFP expression and dead embryos with high GFP expression (**Figure 1C**). *mrp-5* encodes an ABC transporter belonging to the multi-drug resistance protein (MRP) family (Korolnek et al., 2014). MRP proteins are highly similar and the *C. elegans* genome encodes nine family members (www.wormbase.org). To test for specificity, we retested eight of the nine members of this family for which RNAi clones were available, and found that only *mrp-5* RNAi caused activation of GFP expression in the F1 generation (**Figure 1D**). We then titrated vitamin B12 from 6.4 nM to 6.4 mM and found that GFP expression remains activated in *mrp-5* RNAi embryos even under very high vitamin B12 conditions (**Figure S1A**).

MRP-5 is expressed in the intestine (Korolnek et al., 2014). If *mrp-5* indeed encodes a transporter that exports vitamin B12 from the *C. elegans* intestine to support embryonic development, this would lead to the prediction that higher levels of vitamin B12 should be retained in the intestine when the transporter is perturbed. To test this prediction, we devised a highly sensitive experimental setup by using (1) a single copy *Pacdh-1∷GFP∷H2B* strain that expresses GFP in the nucleus, enabling easier monitoring of differences in GFP levels, and (2) an RNAi-compatible strain of *E. coli* OP50 (Xiao et al., 2015), because animals fed this diet express higher levels of GFP than those fed *E. coli* HT115 (**Figure S1B**). Importantly, RNAi of *mrp-5* is equally efficient in *E. coli* HT115 and *E. coli* OP50 as measure by RT-qPCR (**Figure S1C**). We found that much lower concentrations of vitamin B12 were sufficient to repress intestinal GFP in *mrp-5* RNAi animals compared to vector control animals (**Figures 1E** and **1F**). This finding suggests that vitamin B12 is retained in the intestine when *mrp-5* is knocked down. Together these findings indicate that *mrp-5* encodes a transporter that exports vitamin B12 from the intestine of the mother to support embryonic development of her offspring.

### Injection of Vitamin B12 into the Gonad Rescues Embryonic Lethality Caused by Loss of *mrp-5*

We predicted that direct supplementation of vitamin B12 into the gonad should bypass the requirement for vitamin B12 export from the intestine and rescue embryonic lethality caused by perturbation of *mrp-5*. To test this prediction, we injected either vitamin B12 or water (vehicle) as a control into the gonad of *Pacdh-1∷GFP* animals that were exposed to *mrp-5* or vector control RNAi. If the vitamin B12 injection into the gonad bypassed MRP-5-dependent export, we expected to observe live animals with low GFP expression, while injecting water would lead to dead embryos with high GFP expression (**Figure 2A**). For this experiment, we used the *E. coli* OP50 RNAi strains we generated (see also **Figure 1E**) because transgenic *Pacdh-1∷GFP* embryos express high levels of GFP, while they express low GFP levels when fed *E. coli* HT115 bacteria (**Figure S1**). Using *E. coli* OP50 thus enabled us to observe a decrease in GFP expression while using *E. coli* HT115 would not.

**Figure 2.**
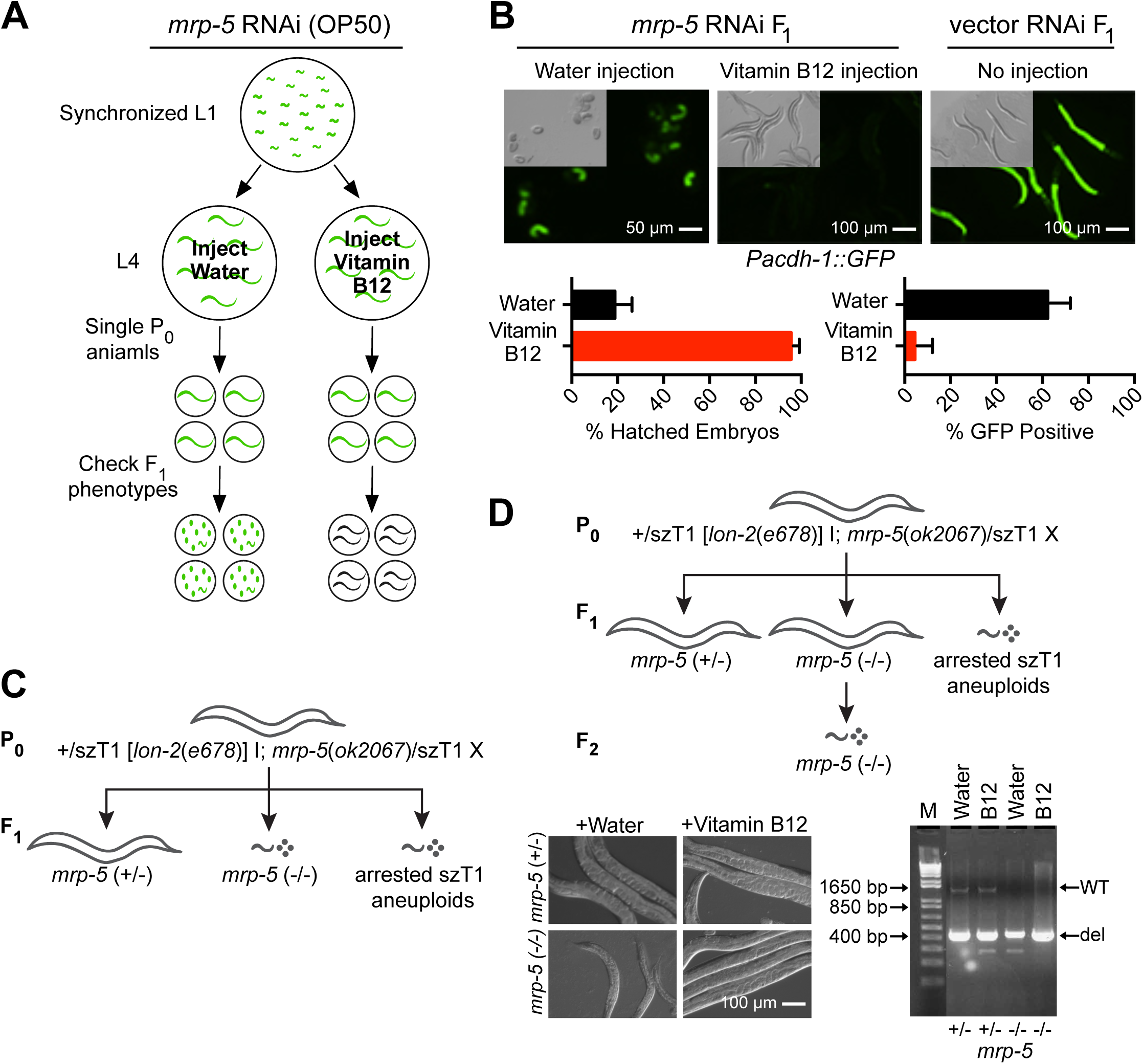
Injection of Vitamin B12 into the Gonad of *mrp-5* deficient Mothers Rescues Lethality in her Offspring. (A) Experimental design. Briefly, mothers were injected with either 3.2 mM vitamin B12 or with water (vehicle) control into the gonad of *Pacdh-1*∷GFP animals that were exposed to *mrp-5* or vector control RNAi. If the vitamin B12 injection rescues embryonic lethality, live offspring will show low GFP expression, while water injected animals will show dead embryos with high GFP expression. (B) Top: DIC and fluorescence microscopy images of embryos from *Pacdh-1∷GFP* mothers subjected to *mrp-5* RNAi in *E. coli* OP50 injected into the gonad with water (left panel) or with 3.2 mM vitamin B12 (middle panel). Animals treated with vector control RNAi without injection are shown as a control (right panel). Scale bars, 50 μm for embryos and 100 μm for adults. Bottom: Quantification of lethality and GFP expression in embryos from *Pacdh-1∷GFP* animals described above. (C) Diagram of genotypes and phenotypes of the offspring of genetically balanced *mrp-5* (+/-) mutant animals. All P0 adult animals are heterozygous for *mrp-5*. F1 larval arrest or embryonic lethality occurs in *mrp-5* (-/-) or szT1 aneuploid animals. (D) Top: Diagram of genotypes and phenotypes of viable offspring from genetically balanced *mrp-5* (+/-) mutant animals after vitamin B12 injection. All P0 adult animals are heterozygous for *mrp-5*. Live F1 animals can either be heterozygous or homozygous for the *mrp-5*(ok2067) deletion allele. F2 larval arrest or embryonic lethality occurs in all offspring of the F2 generation of *mrp-5* (-/-) animals. Bottom left: DIC microscopy images of *mrp-5* (+/-) and *mrp-5* (-/-) animals with or without vitamin B12 injection. Scale bar, 100 μm. Bottom right: Agarose gel image of the genotyping results of heterozygous and homozygous *mrp-5* adult animals with and without injection of vitamin B12.

Injection of vitamin B12 into the gonad of *mrp-5* RNAi animals fully and specifically rescued embryonic viability of their offspring (**Figure 2B**). In addition, in *Pacdh-1∷GFP* transgenic animals exposed to *mrp-5* RNAi, injecting vitamin B12 into the gonad resulted in developed larvae with low levels of GFP, while injection of water resulted in dead embryos with high GFP expression (**Figure 2B**). We recapitulated these findings with *mrp-5(ok2067*) deletion mutant animals. Since *mrp-5* is an essential gene, the *mrp-5* deletion allele is maintained genetically balanced with +/svT1[*lon-2*(e678)] (**Figure 2C**). Heterozygous *mrp-5* (+/-) mothers injected with vitamin B12 into their gonad produced *mrp-5* (-/-) offspring that survived to the adult stage, while animals injected with vehicle control produced *mrp-5* (-/-) animals that arrested at the L1-L2 stage (**Figure 2D**). Importantly, offspring of the rescued F1 *mrp-5* (-/-) animals again exhibited larval arrest, suggesting that in the absence of *mrp-5* vitamin B12 cannot be passed on to the next (F2) generation. Altogether, these findings show that vitamin B12 injected into the gonad of a *mrp-5* (+/-) mother can rescue embryonic lethality in her offspring.

### Heme Rescues Embryonic Lethality in *mrp-5* RNAi Animals

A previous study proposed that *mrp-5* encodes a heme transporter (Korolnek et al., 2014). This finding was supported by the observation that embryonic lethality in *mrp-5* RNAi animals could be rescued by feeding the animals high doses of heme (Korolnek et al., 2014). How could either feeding of heme or injection of vitamin B12 rescue embryonic lethality of *mrp-5* RNAi or mutant animals? ABC transporters are known to be capable of transporting numerous similar types of molecules (Lee and Rosenbaum, 2017; Locher, 2016; Slot et al., 2011). Heme and vitamin B12 share some structural similarities; heme is an iron-containing porphyrin, while vitamin B12 is related to porphyrins and contains cobalt as a metal ion (**Figure 3A**). Thus, it is possible that MRP-5 transports both molecules. However, if lack of heme *and* vitamin B12 explains embryonic lethality of *mrp-5* RNAi animals, one would expect that neither cofactor alone would be sufficient for rescue. Our observation that embryonic lethality can be rescued by injection of vitamin B12 into the gonad indicates that lack of vitamin B12 rather than heme may be the cause of embryonic lethality when MRP-5 function is perturbed. To further examine the effects of heme on animal development, we repeated the experiment published in the previous study (Korolnek et al., 2014) and confirmed that feeding 500 μM heme can rescue embryonic lethality in the offspring of *mrp-5* RNAi animals and allowed some animals to develop to L2 and later stages (**Figure 3B**). Interestingly, however, such a high dose of heme was toxic to vector control animals, arresting most animals at the L1-L2 stage (**Figure 3B**). Importantly, one can argue that feeding heme does not actually test for transport because the cofactor would be stuck in the intestine when *mrp-5* is perturbed, which would thus not result in the rescue of embryonic development in the offspring.

**Figure 3.**
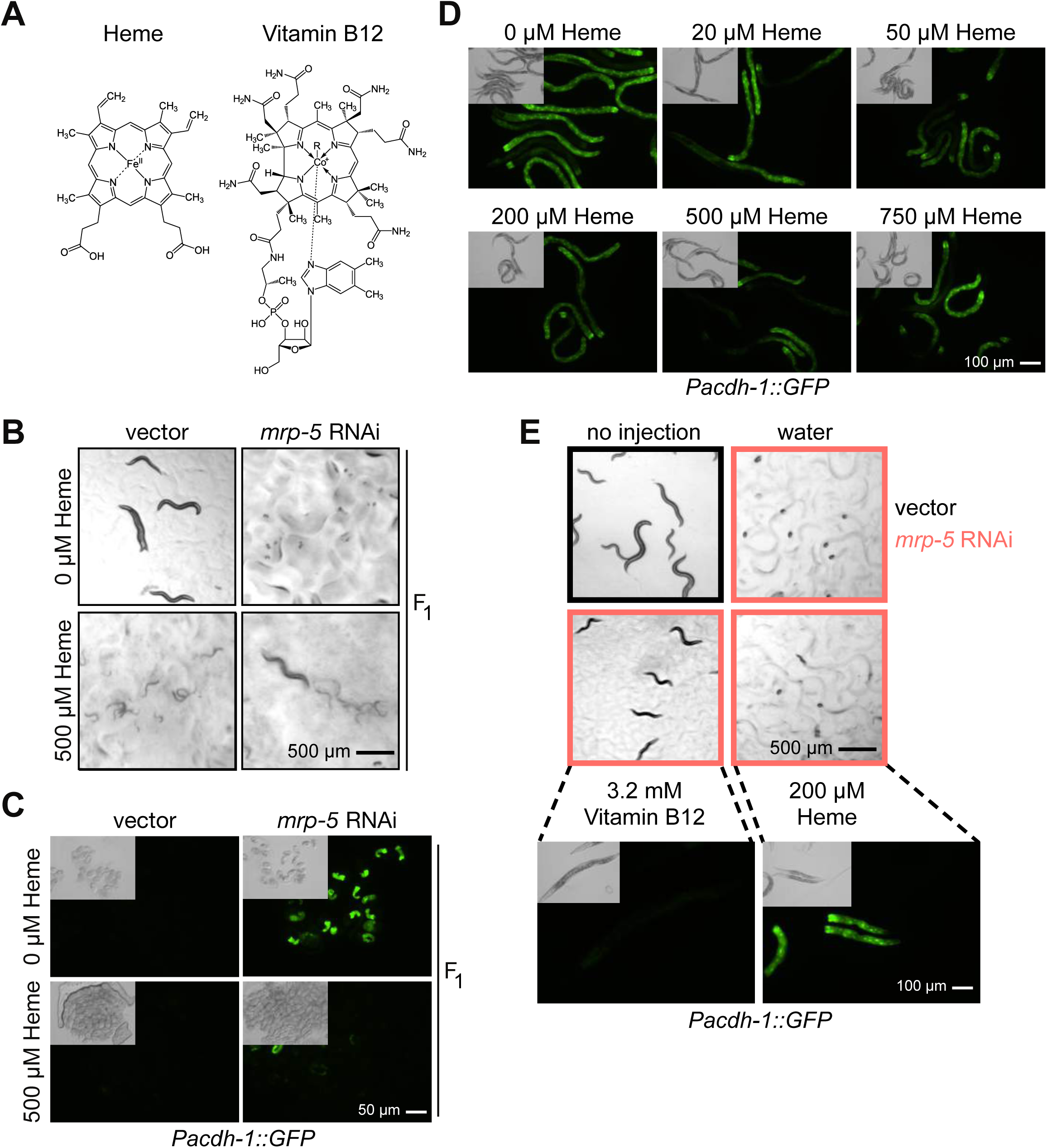
High Concentrations of Heme Rescue Embryonic Lethality in *mrp-5* RNAi Animals but Do Not Affect *acdh-1* Promoter Activity. (A) Heme and vitamin B12 are porphyrin or porphyrin-like molecules with structural similarities. (B) Bright field images of feeding 500 μM heme to *mrp-5* RNAi-treated mothers on *E. coli* HT115 rescues viability in her offspring and allows them to develop to the L2 stage and later, but causes L1 larval arrest and toxicity in vector-treated mothers. Scale bar, 500 μm. (C) Fluorescence and DIC microscopy images show that feeding 500 μM heme to *mrp-5* RNAi-treated mothers on *E. coli* HT115 reduces *Pacdh-1∷GFP* reporter activity in embryos, but not in embryos of vector control mothers. Scale bar, 50 μm. (D) Fluorescence and DIC microscopy images demonstrate that feeding heme to mothers with *E. coli* OP50 does not affect *Pacdh-1∷GFP* reporter activity. Scale bar, 100 μm. (E) Bright field (top) and fluorescence and DIC microscopy images (bottom) shows injection of 200 μM heme into *mrp-5* RNAi-treated mothers with *E. coli* HT115 does not rescue embryonic lethality or larval arrest in her offspring, while injection of vitamin B12 leads to viable offspring and repression of the *acdh-1* promoter. Scale bar, 500 μm for bright field and 100 μm for fluorescence microscopy.

One explanation for the rescue of embryonic development by feeding heme is that this increases vitamin B12 passage through intestinal membranes. Therefore, we next asked whether heme affects the vitamin B12-responsive *Pacdh-1* reporter. We supplemented increasing concentrations of heme and found that the reporter was unaffected by heme in adult animals (**Figure 3D**). As noted above, embryos from *Pacdh-1∷GFP* mothers treated with *mrp-5* RNAi without supplemented heme expressed high levels of GFP, suggesting that vitamin B12 cannot enter the embryos and repress the reporter. Importantly, however, GFP levels were greatly reduced in embryos from *Pacdh-1∷GFP/mrp-5* RNAi animals supplemented with 500 μM heme (**Figure 3C**). These findings suggest that supplying a high dose of heme facilitates export of vitamin B12 from the intestine and, therefore, that the cause for embryonic lethality is lack of vitamin B12 and not heme. Indeed, we found that while injection of vitamin B12 into the gonad rescued embryonic development in *Pacdh-1∷GFP/mrp-5* RNAi animals, direct injection of 200 μM heme did not rescue. Further, the arrested L1 larvae from *Pacdh-1∷GFP*/*mrp-5* RNAi animals injected with 200 μM heme expressed high levels of GFP indicating low vitamin B12 levels (**Figure 3E**).

### Methionine and S-Adenosyl Methionine Content is Reduced in Offspring from *mrp-5* RNAi Mothers

Although vitamin B12 is a cofactor for two different metabolic enzymes (**Figure 1A**), our previous studies demonstrated that the developmental delay caused by vitamin B12 deficiency is mainly due to its function as a cofactor for the methionine synthase enzyme METR-1, which generates the methyl donor SAM from methionine (Watson et al., 2014). To determine if the lack of vitamin B12 or the lack of heme in *mrp-5* deficient animals results in defects in the SAM cycle, we measured the levels of methionine and SAM in embryos from mothers treated with *mrp-5* or vector control RNAi. We reasoned that differences in methionine and SAM content would be most clear in the presence of supplemented vitamin B12 because the *E. coli* diet is naturally low in this cofactor. Indeed, we detected a reduced methionine and SAM content in embryos from mothers treated with *mrp-5* RNAi relative to control animals in the presence of vitamin B12 (**Figures 4A** and **4B**). Further, SAM levels could be restored to wild type levels by treating the animals with 500 μM heme. These results suggest that a defect in the SAM cycle caused by the lack of vitamin B12 in *mrp-5* deficient embryos contributes to embryonic lethality.

**Figure 4.**
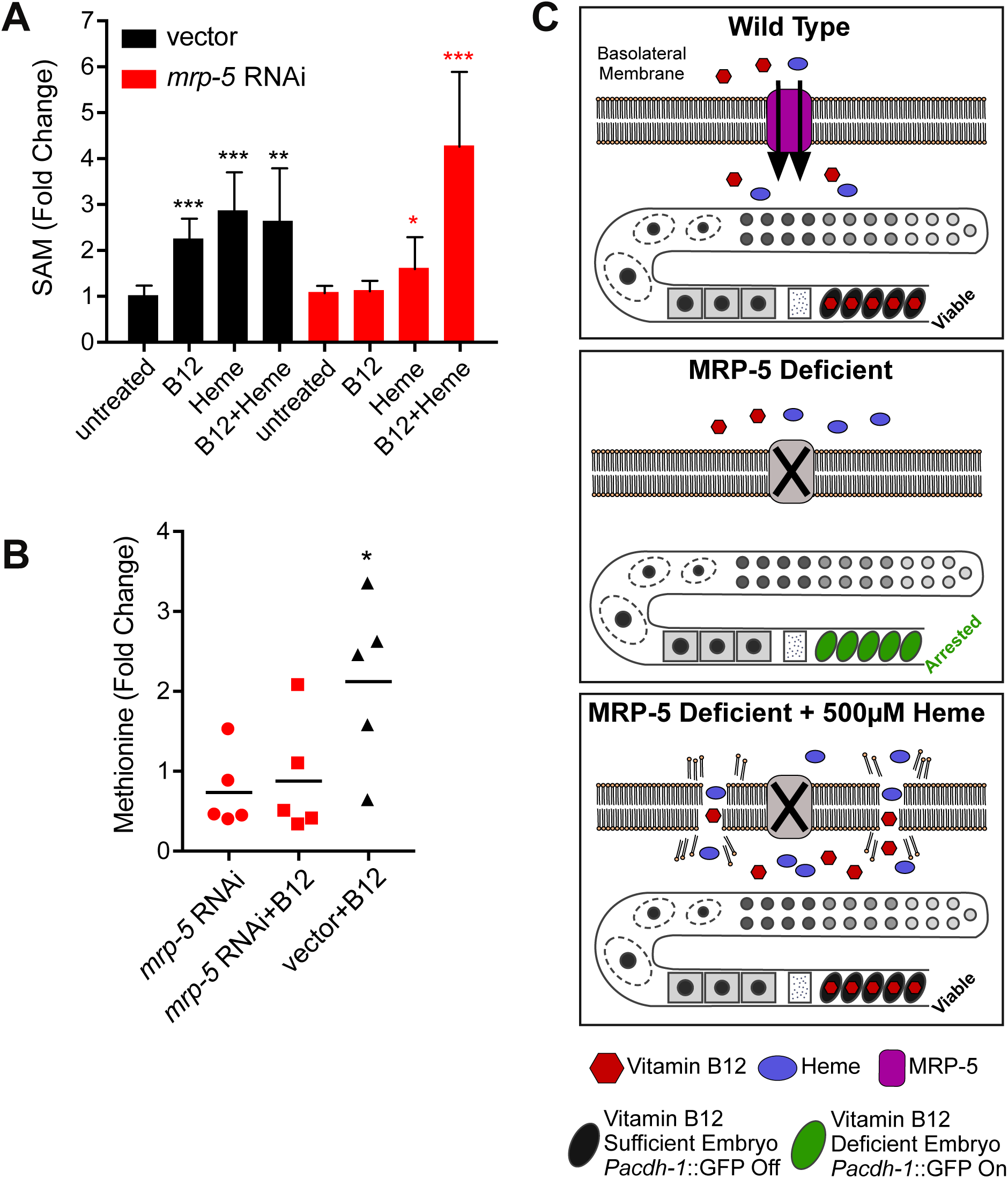
Methionine and S-Adenosyl Methionine Content is Reduced in Offspring from Mothers Treated with *mrp-5* RNAi. (A) SAM content was measured in the embryos of vector or *mrp-5* RNAi-treated mothers on *E. coli* HT115. SAM content was greatly reduced in *mrp-5* deficient embryos compared to vector control. *** p < 0.0001, ** p < 0.001, * p < 0.05 relative to untreated embryos. (Student’s t-test). (B) Methionine content was measured by GC-MS of vector control or *mrp-5* RNAi-treated mothers fed *E. coli* HT115 with and without 64 nM vitamin B12 supplementation. Each dot represents a biological replicate, and values are relative to vector RNAi without vitamin B12 supplementation. (C) Model of MRP-5 function. In wild type animals both vitamin B12 and heme are exported from the intestine by MRP-5. Vitamin B12 can enter the embryo and repress the *acdh-1* reporter (top). In the absence of MRP-5 neither vitamin B12 nor heme can cross the intestinal membrane, the embryos arrest, and the *acdh-1* reporter remains active (middle). The presence of 500 μM heme causes membrane permeability in *mrp-5*-deficient mothers leading to vitamin B12 crossing the intestine membrane, entering the embryo to rescue lethality, and repressing the *acdh-1* promoter (bottom).

## DISCUSSION

We identified MRP-5 as a candidate transporter that exports vitamin B12 from the *C. elegans* intestine of the mother to support the development of her offspring. MRP-5 is a member of the *C. elegans* ABC transporter family, and these transporters are known to function as exporters of metabolites and drugs (Locher, 2016; Sheps et al., 2004; Slot et al., 2011). MRP-5 has previously been reported to export heme in *C. elegans* (Korolnek et al., 2014). *C. elegans* can synthesize neither vitamin B12 nor heme and depends on dietary supplementation of both cofactors. Loss or reduction of *mrp-5* by RNAi knockdown or by genetic mutation results in embryonic or larval lethality. As reported previously, we confirm that this lethality can be rescued by feeding the mother high concentrations of heme. However, it can also be rescued by injecting vitamin B12 directly into the gonad, bypassing the need for import from the intestine. Wild type *C. elegans* require only low doses of heme (20 μM) (Rao et al., 2005) and high doses of heme are toxic to many systems (Chiabrando et al., 2014). Indeed, supplementing wild type animals with a dose of 500 μM heme was toxic to *C. elegans* as well (**Figure 3B**). One mechanism of heme toxicity is by enhanced membrane permeability (Chiabrando et al., 2014). Therefore, we propose that supplementing *mrp-5* RNAi animals with high concentrations of heme leads to increased intestinal membrane permeability, enabling both heme and vitamin B12 to pass through without the need for a dedicated transporter (**Figure 4C**).

More than 15 human genes involved in vitamin B12 processing have been identified (Nielsen et al., 2012), and four of these encode transporters. One of these is the ABC transporter MRP-1 (or ABCC1), which is localized to intestinal epithelial cells. ABC transporters in human and *C. elegans* are highly similar in sequence and structure. Therefore, we propose that *C. elegans* MRP-5 may be a functional ortholog of human MRP1 to export vitamin B12 from the intestine to surrounding tissues.

## ACKNOWLEDGEMENTS

We thank members of the Walhout lab and Amy Walker for discussion and critical reading of the manuscript. We thank Amy Holdorf for manuscript editing and Shawn Xu for the *E. coli* OP50 RNAi strain. This work was supported by a grant from the National Institutes of Health (DK068429) to A.J.M.W. Some bacterial and nematode strains used in this work were provided by the CGC, which is funded by the NIH Office or Research Infrastructure Programs (P40 OD010440).

## AUTHOR CONTRIBUTIONS

H.N. and A.J.M.W. conceived the study. H.N. performed all the experiments. G.G. generated the single copy *Pacdh-1∷H2B∷GFP* strain. O.P. performed the GC-MS analysis. H.N. and A.J.M.W. wrote the manuscript.

## STAR ★ METHODS

### KEY RESOURCES TABLE

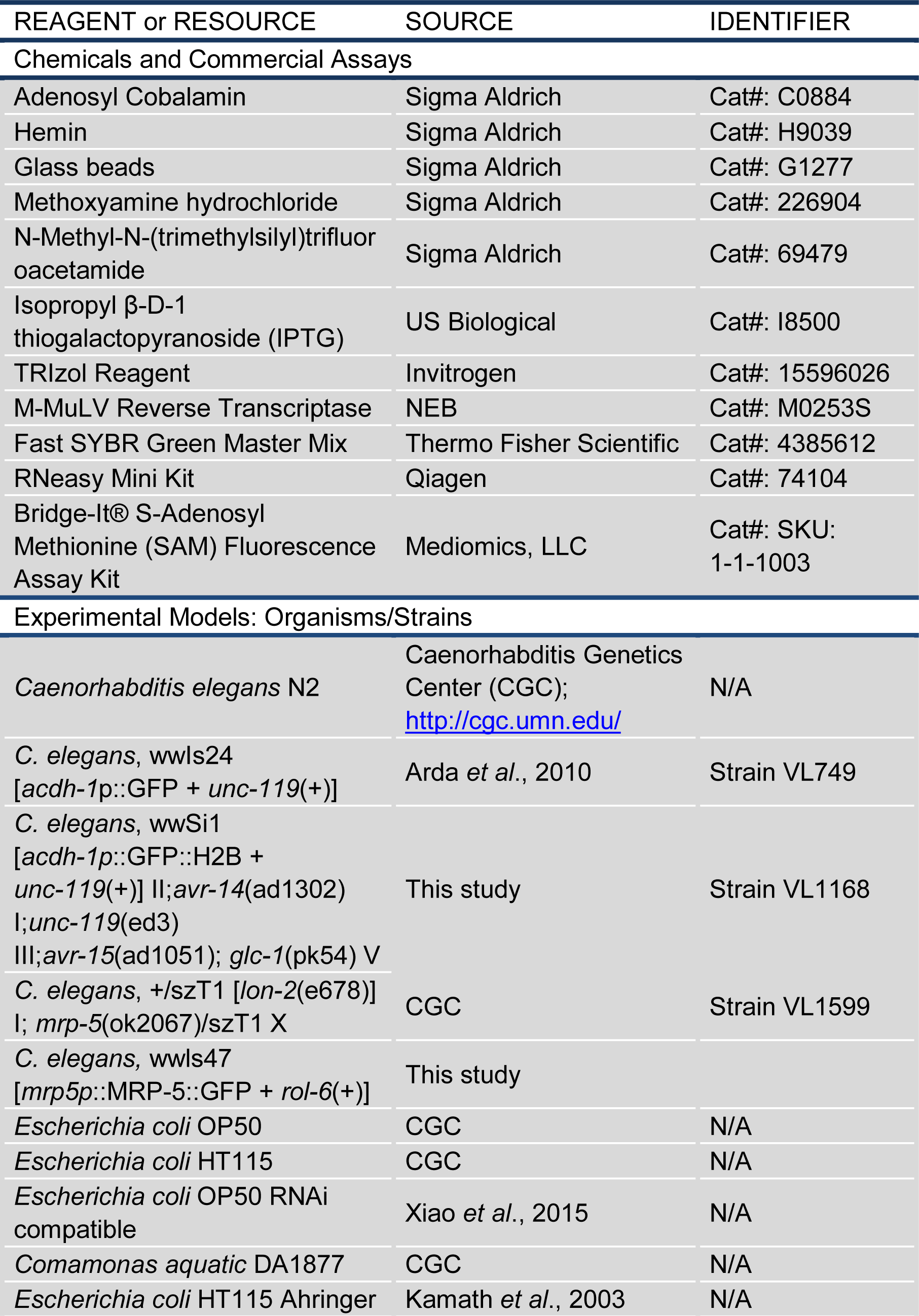
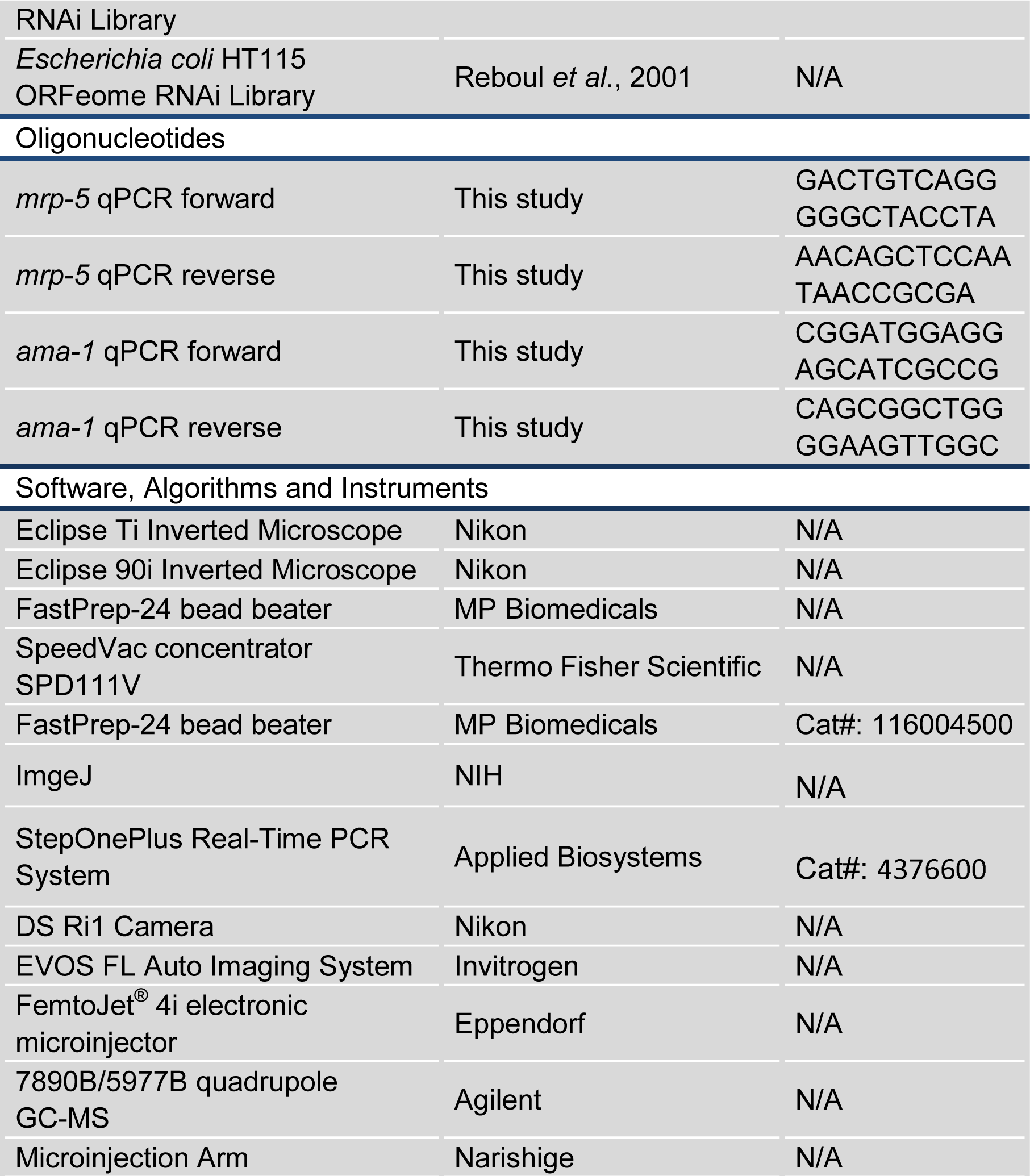

## EXPERIMENTAL PROCEDURES

### *C. elegans* Strains

All *C. elegans* strains were maintained at standard laboratory conditions as described (Brenner, 1974), and strain N2 was used as wild type. Construction of VL749 (wwIs24 [*Pacdh-1∷GFP* + unc-119(+)]) strain has been previously described (Arda et al., 2010). Strain VL1168 (*wwSi1[Pacdh-1∷GFP∷H2B unc-119(+)] II;avr-14(ad1302) I; unc-119(ed3) III; avr-15(ad1051); glc-1(pk54) V*) was generated by *mos1*-mediated single copy insertion (Frokjaer-Jensen et al., 2008). Strain wwls47 [*mrp5p*∷MRP-5∷GFP + *rol-6*(+)] was generated by fusion PCR (www.wormbook.org). The *mrp-5* mutant strain VC1599 (+/szT1 [*lon-2*(e678)] I; *mrp-5*(ok2067)/szT1 X) was obtained from the *C. elegans* Genetics Center (CGC). Bacterial strains *E. coli* OP50, HT115 and *Comamonas aquatica* DA1877 were obtained from CGC. The *E. coli* OP50 RNAi compatible bacterial strain was generously provided by the Xu lab (Xiao et al., 2015).

### RNAi Screen

The Kyoto Encyclopedia of Genes and Genomes (KEGG) database was used to predict *C. elegans* ABC transporters and solute carrier transporters (Kanehisa et al., 2015). The transporter RNAi mini-library was generated by selecting 215 clones from both the ORFome and Ahringer RNAi libraries (Kamath et al., 2003; Rual et al., 2004). RNAi screening was carried out on 24-well Nematode Growth Medium (NGM) agar plates containing 2 mM isopropyl β-D-1-thiogalactopyranoside (IPTG), 50 μg/ml ampicillin with or without 64 nM Ado-Cbl (vitamin B12). *Pacdh-1*∷GFP animals were grown on regular NGM plates with *E. coli* OP50 without Ado-Cbl supplementation for several generations before harvesting embryos. Embryos were incubated in M9 buffer overnight to obtain synchronized L1 animals. Approximately 15 L1 animals were added to each well containing individual RNAi clones. Animal phenotypes were observed after 80 hours of incubation, when the F1 generation was at the young adult stage.

### Quantitation of Fluorescence

The animal body was outlined and fluorescence intensity of the animal was determined using ImageJ (NIH). At least five animals per condition were analyzed, and the mean fluorescence per animals was determined.

### *C. elegans* qRT-PCR Experiments

Animals were synchronized by L1 arrest and grown on ampicillin and IPTG plates seeded with either *E. coli* HT115 (vector RNAi and *mrp-5* RNAi) or *E. coli* OP50 (vector RNAi and *mrp-5* RNAi). Approximately 1000 adult animals were harvested for each condition, in biological duplicate. Animals were washed in M9 buffer, and total RNA was isolated using TRIzol (Invitrogen) followed by DNAseI treatment and cleanup using Qiagen RNeasy columns. cDNA was generated from 1 μg RNA using random primer and M-MuLV reverse transcriptase (NEB). qPCR was performed in technical duplicates per gene per condition using the Applied Biosystems StepOnePlus Real-Time PCR system and Fast SYBR Green Master Mix (Thermo Fisher Scientific). Relative transcript abundance was determined using the ΔΔCt method, and normalized to averaged *ama-1* mRNA expression levels.

### Vitamin B12 Injections

Animals were treated with *mrp-5* RNAi generated in an *E. coli* OP50 RNAi compatible strain. Vitamin B12 (3.2 mM) was injected into the gonad of L4 animals using a Narishige microinjection arm attached to the body of a Nikon Eclipse Ti inverted microscope. Sterile water was injected into the control group. After injection, animals were singled and treated with the same RNAi (*mrp-5* or vector). Phenotypes were scored after two days. *mrp-5*(ok2067) heterozygotes were singled at the L4 stage. Vitamin B12 was injected as above. After injection, animals were singled and the phenotype was scored after five days.

### S-Adenosyl Methionine (SAM) Measurement

SAM was measured using a Bridge-It S-Adenosyl Methionine Fluorescence assay kit purchased from Mediomics following the manufacturer’s recommendations.

### Heme Solution Preparation

10 mM Hemin solution was prepared as described (Hickman and Winston, 2007). 6.5 mg/ml Hemin (Sigma-Aldrich) was dissolved in 0.1 M NaOH and incubated at 37°C for 1 hour. 1 M Tris, pH 7.5, was then added to a final concentration of 0.1 M. The pH was adjust using HCl, and the hemin solution was stored at 4°C, protected from light and used within two days.

### Relative Quantification of Methionine using GC-MS

Approximately 150,000 embryos were homogenized with 0.5 ml of 200-300 μm acid-washed glass beads (Sigma-Aldrich) in 1 ml 80% methanol using FastPrep-24 bead beater (MP Biomedicals), with intermittent cooling in dry ice/ethanol bath. Samples were then extracted on dry ice for 15 min., centrifuged for 10 min. at 20,000xg and 250 μl of supernatant were dried under vacuum using a SpeedVac concentrator SPD111V (Thermo Fisher Scientific). Derivatization of dried samples was performed first with 20 μl of 20 mg/ml methoxyamine hydrochloride (Sigma-Aldrich) in pyridine at 37°C for 90 minutes, followed by addition of 50 μl of N-Methyl-N-(trimethylsilyl)trifluoroacetamide (Sigma-Aldrich) and incubation for 3 hours at 37°C. The derivitazation reaction was completed by incubation for 5 hours at room temperature. Measurements were performed on an Agilent 7890B/5977B quadrupole GC-MS equipped with 30 m x 0.25 mm x 0.25 μm HP-5MS UI capillary column. The inlet temperature was set to 230°C, transfer line was at 280°C, MS source and quadrupole were at 230°C and 150°C respectively. Trimethylsilyl derivative of methionine was quantified as a 176 m/z ion with two qualifier ions 128 and 293 m/z. Relative quantification of peak areas was done using samples within a linear response range, after mean normalization to total metabolites and blank subtraction.

## SUPPLEMENTAL INFORMATION

Supplemental information includes two figures.

**Figure S1**.

(A) Fluorescence and DIC microscopy images of embryos from *Pacdh-1∷GFP* mothers subjected to vector control RNAi or *mrp-5* RNAi and supplemented with increasing concentrations of vitamin B12. In the upper row, the inset box in the lower right indicates 15 times the exposure time. Scale bar, 50 μm.

(B) Fluorescence and DIC microscopy images of *Pacdh-1∷GFP* animals fed *E. coli* OP50, *E. coli* HT115 or *C. aquatica* DA1877 at P0 and F1 generations. Scale bars, 50 μm for embryos and 100 μm for adults.

(C) RT-qPCR of *mrp-5* showing similar RNAi efficiency in *E. coli* HT115 and *E. coli* OP50-fed animals relative to vector control.

**Figure S2**.

Fluorescence (top), DIC (middle), and overlay (bottom) microscopy images of GFP expression in *Pmrp-5∷mrp-5∷GFP* transgenic animals. The white arrowheads in the middle panels indicate intestinal cells.

## REFERENCES

Arda, H.E., Taubert, S., Conine, C., Tsuda, B., Van Gilst, M.R., Sequerra, R., Doucette-Stam, L., Yamamoto, K.R., and Walhout, A.J.M. (2010). Functional modularity of nuclear hormone receptors in a C. elegans gene regulatory network. Mol. Syst. Biol. 6, 367.

Bito, T., Matsunaga, Y., Yabuta, Y., Kawano, T., and Watanabe, F. (2013). Vitamin B12 deficiency in Caenorhabditis elegans results in loss of fertility, extended life cycle, and reduced lifespan. FEBS Open Bio. 3, 112–117.

Brenner, S. (1974). The genetics of Caenorhabditis elegans. Genetics 77, 71–94.

Carrillo-Carrasco, N., and Venditti, C. (1993). Propionic Acidemia. In GeneReviews, R.A. Pagon, M.P. Adam, H.H. Ardinger, T.D. Bird, C.R. Dolan, C.T. Fong, R.J.H. Smith, and K. Stephens, eds. (Seattle (WA)).

Chiabrando, D., Vinchi, F., Fiorito, V., Mercurio, S., and Tolosano, E. (2014). Heme in pathophysiology: a matter of scavenging, metabolism and trafficking across cell membranes. Front. Pharmacol. 5, 61.

Czeizel, A.E., and Dudas, I. (1992). Prevention of the first occurrence of neural-tube defects by periconceptional vitamin supplementation. N. Engl. J. Med. 327, 1832–1835.

Dror, D.K., and Allen, L.H. (2008). Effect of vitamin B12 deficiency on neurodevelopment in infants: current knowledge and possible mechanisms. Nutr. Rev. 66, 250–255.

Fall, C.H., Yajnik, C.S., Rao, S., Davies, A.A., Brown, N., and Farrant, H.J. (2003). Micronutrients and fetal growth. J. Nutr. 133, 1747S–1756S.

Frokjaer-Jensen, C., Davis, M.W., Hopkins, C.E., Newman, B.J., Thummel, J.M., Olesen, S.P., Grunnet, M., and Jorgensen, E.M. (2008). Single-copy insertion of transgenes in Caenorhabditis elegans. Nat. Genet. 40, 1375–1383.

Hickman, M.J., and Winston, F. (2007). Heme levels switch the function of Hap1 of Saccharomyces cerevisiae between transcriptional activator and transcriptional repressor. Mol. Cell. Biol. 27, 7414–7424.

Kamath, R.S., Fraser, A.G., Dong, Y., Poulin, G., Durbin, R., Gotta, M., Kanapin, A., Le Bot, N., Moreno, S., Sohrmann, M., et al. (2003). Systematic functional analysis of the Caenorhabditis elegans genome using RNAi. Nature 421, 231–237.

Kanehisa, M., Sato, Y., Kawashima, M., Furumichi, M., and Tanabe, M. (2016). KEGG as a reference resource for gene and protein annotation. Nucleic Acids Res. 44, 457–462.

Korolnek, T., Zhang, J., Beardsley, S., Scheffer, G.L., and Hamza, I. (2014). Control of metazoan heme homeostasis by a conserved multidrug resistance protein. Cell Metab. 19, 1008–1019.

Lee, J.Y., and Rosenbaum, D.M. (2017). Transporters Revealed. Cell 168, 951–953.

Locher, K.P. (2016). Mechanistic diversity in ATP-binding cassette (ABC) transporters. Nat. Struct. Mol. Biol. 23, 487–493.

MacNeil, L.T., Watson, E., Arda, H.E., Zhu, L.J., and Walhout, A.J.M. (2013). Diet-induced developmental acceleration independent of TOR and insulin in C. elegans. Cell 153, 240–252.

Martens, J.H., Barg, H., Warren, M.J., and Jahn, D. (2002). Microbial production of vitamin B12. Appl. Microbiol. Biotechnol. 58, 275–285.

Moreno-Garcia, M.A., Rosenblatt, D.S., and Jerome-Majewska, L.A. (2013). Vitamin B(12) metabolism during pregnancy and in embryonic mouse models. Nutrients 5, 3531–3550.

Nielsen, M.J., Rasmussen, M.R., Andersen, C.B., Nexo, E., and Moestrup, S.K. (2012). Vitamin B12 transport from food to the body’s cells—a sophisticated, multistep pathway. Nat. Rev. Gastroenterol. Hepatol. 9, 345–354.

Owens, S., and Fall, C.H. (2008). Consequences of poor maternal micronutrition before and during early pregnancy. Trans. R. Soc. Trop. Med. Hyg. 102, 103–104.

Rao, A.U., Carta, L.K., Lesuisse, E., and Hamza, I. (2005). Lack of heme synthesis in a free-living eukaryote. Proc. Natl. Acad. Sci. USA 102, 4270–4275.

Rosenblatt, D.S., and Cooper, B.A. (1990). Inherited disorders of vitamin B12 utilization. Bioessays 12, 331–334.

Rosenblatt, D.S., and Whitehead, V.M. (1999). Cobalamin and folate deficiency: acquired and hereditary disorders in children. Semin. Hematol. 36, 19–34.

Rual, J.-F., Ceron, J., Koreth, J., Hao, T., Nicot, A.-S., Hirozane-Kishikawa, T., Vandenhaute, J., Orkin, S.H., Hill, D.E., van den Heuvel, S., et al. (2004). Toward improving Caenorhabditis elegans phenome mapping with an ORFeome-based RNAi library. Genome Res. 14, 2162–2168.

Sheps, J.A., Ralph, S., Zhao, Z., Baillie, D.L., and Ling, V. (2004). The ABC transporter gene family of Caenorhabditis elegans has implications for the evolutionary dynamics of multidrug resistance in eukaryotes. Genome Biol. 5, R15.

Skjaerven, K.H., Jakt, L.M., Dahl, J.A., Espe, M., Aanes, H., Hamre, K., and Fernandes, J.M. (2016). Parental vitamin deficiency affects the embryonic gene expression of immune-, lipid transport-and apolipoprotein genes. Sci. Rep. 6, 34535.

Slot, A.J., Molinski, S.V., and Cole, S.P. (2011). Mammalian multidrug-resistance proteins (MRPs). Essays Biochem. 50, 179–207.

Viswanathan, M., Treiman, K.A., Kish-Doto, J., Middleton, J.C., Coker-Schwimmer, E.J., and Nicholson, W.K. (2017). Folic Acid Supplementation for the Prevention of Neural Tube Defects: An Updated Evidence Report and Systematic Review for the US Preventive Services Task Force. JAMA 317, 190–203.

Watson, E., MacNeil, L.T., Arda, H.E., Zhu, L.J., and Walhout, A.J.M. (2013). Integration of metabolic and gene regulatory networks modulates the C. elegans dietary response. Cell 153, 253–266.

Watson, E., MacNeil, L.T., Ritter, A.D., Yilmaz, L.S., Rosebrock, A.P., Caudy, A.A., and Walhout, A.J.M. (2014). Interspecies systems biology uncovers metabolites affecting C. elegans gene expression and life history traits. Cell 156, 759–770.

Watson, E., Olin-Sandoval, V., Hoy, M.J., Li, C.-H., Louisse, T., Yao, V., Mori, A., Holdorf, A.D., Troyanskaya, O.G., Ralser, M., et al. (2016). Metabolic network rewiring of propionate flux compensates vitamin B12 deficiency in C. elegans. Elife 5, pii: e17670.

Xiao, R., Chun, L., Ronan, E.A., Friedman, D.I., Liu, J., and Xu, X.Z. (2015). RNAi interrogation of dietary modulation of development, metabolism, behavior, and aging in C. elegans. Cell. Rep. 11, 1123–1133.

Yilmaz, L.S., and Walhout, A.J.M. (2014). Worms, bacteria and micronutrients: an elegant model of our diet. Trends. Genet. 30, 496–503.

